# MYC is sufficient to generate mid-life high-grade serous ovarian and uterine serous carcinomas in a p53-R270H mouse model

**DOI:** 10.1101/2024.01.24.576924

**Authors:** Alexandra Blackman, Amy C Rees, Robert R Bowers, Christian M Jones, Silvia G Vaena, Madison A Clark, Shelby Carter, Evan D Villamor, Della Evans, Anthony J Emanuel, George Fullbright, David T Long, Laura Spruill, Martin J Romeo, Kristi L Helke, Joe R Delaney

**Author notes:** Equal contributions (Alexandra Blackman, Amy C Rees, Robert R Bowers, Christian M Jones). Correspondence to: JR Delaney, Department of Biochemistry and Molecular Biology, Medical University of South Carolina, 173 Ashley Ave, Charleston, SC 29425, USA.

## Abstract

Genetically engineered mouse models (GEMM) have fundamentally changed how ovarian cancer etiology, early detection, and treatment is understood. However, previous GEMMs of high-grade serous ovarian cancer (HGSOC) have had to utilize genetics rarely or never found in human HGSOC to yield ovarian cancer within the lifespan of a mouse. *MYC*, an oncogene, is amongst the most amplified genes in HGSOC, but it has not previously been utilized to drive HGSOC GEMMs. We coupled *Myc* and dominant negative mutant p53-R270H with a fallopian tube epithelium-specific promoter *Ovgp1* to generate a new GEMM of HGSOC. Female mice developed lethal cancer at an average of 15.1 months. Histopathological examination of mice revealed HGSOC characteristics including nuclear p53 and nuclear MYC in clusters of cells within the fallopian tube epithelium and ovarian surface epithelium. Unexpectedly, nuclear p53 and MYC clustered cell expression was also identified in the uterine luminal epithelium, possibly from intraepithelial metastasis from the fallopian tube epithelium (FTE). Extracted tumor cells exhibited strong loss of heterozygosity at the p53 locus, leaving the mutant allele. Copy number alterations in these cancer cells were prevalent, disrupting a large fraction of genes. Transcriptome profiles most closely matched human HGSOC and serous endometrial cancer. Taken together, these results demonstrate the *Myc* and *Trp53-R270H* transgene was able to recapitulate many phenotypic hallmarks of HGSOC through the utilization of strictly human-mimetic genetic hallmarks of HGSOC. This new mouse model enables further exploration of ovarian cancer pathogenesis, particularly in the 50% of HGSOC which lack homology directed repair mutations. Histological and transcriptomic findings are consistent with the hypothesis that uterine serous cancer may originate from the fallopian tube epithelium.

## Introduction

Gynecologic oncology has only recently been able to take advantage of the targeted therapies generated over the last decade, despite there being great need. Ovarian cancer is the 5th most common cancer death in women and uterine cancer is the 6th. Ovarian cancer deaths are dominated by the high-grade serous ovarian cancer (HGSOC) subtype. Uterine cancer, overall, is less lethal despite higher incidence rates, yet the serous carcinoma subtype accounts for 40% of endometrial cancer deaths [1]. Uterine cancer incidence and mortality has increased due to a disproportionate increase in type 2 or non-endometrioid subtypes, including serous carcinoma [2].

Recent targeted therapies in HGSOC involve PARP inhibitors (PARPi), which best benefit patients with *BRCA1* or *BRCA2* tumor mutations or markers of homologous repair deficiency: half of patients in sum [3]. Uterine cancers have shown promise in large Phase III trials of immunotherapies, particularly in the mismatch repair deficient cases [4, 5], but only a small subset of serous carcinoma patients is mismatch repair deficient. Unfortunately, few targeted drugs have shown clinical benefit for HGSOC with normal homologous repair or in recurrent tumors which restore homologous repair after PARPi therapy. Similarly, serous endometrial cancer has few targetable alterations and thus few drugs in development [6]. Serous endometrial cancer and HGSOC share similar disease etiology, such as post-menopausal disease presentation and peritoneal metastasis, and share similar molecular characteristics, such as low mutation rate, high copy-number alterations, mutation in p53, and few other driver mutations.

Genetically engineered mouse models (GEMM) have fundamentally changed how ovarian cancer etiology and treatment is understood. Phylogenetic assessments of HGSOC compared primary tumors dissected from an ovary and metastases, which demonstrated the earliest mutations often originated in serous tubal intraepithelial carcinoma (STIC) [7]. However, this study and other studies have shown that cells can arise first on the ovarian surface epithelium (OSE) or other origin sites, metastasize to the fallopian tube epithelium (FTE) and develop a STIC, and finally develop into HGSOC [8–10]. A recent pair of GEMMs indicate that OSE-originating cancer cells are phenotypically distinct from fallopian fimbriae-originating cancer cells, despite identical drivers [9]. *Lgr5* was used to drive OSE gene deletion while *Pax8* was used to drive FTE gene deletion. Previous research established the *Ovgp1* model of oviductal transgene expression, where *Ovgp1*-mediated expression control was exclusive to the FTE [11, 12].

These previous mouse models revolutionized the field but have some important limitations. Most models utilized homozygous deletions of multiple tumor suppressors to generate tumors, which is not often, if ever, found in human disease. We previously investigated aneuploidy with or without a heterozygous deletion of the autophagy gene *Becn1* in the *Amhr2*-SV40-T-antigen model [13, 14]. These viral proteins are known to disrupt p53 function and RB1 function, amongst other proteins [15, 16]. However, few aneuploid events were observed relative to human disease [14], similar to other GEMM findings [17]. Previous HGSOC mouse models utilize *BRCA* deletions, *PTEN* deletions, or viral oncogenes. The field is left without a genetically representative model for HGSOC lacking *BRCA1/2* or other homologous repair alterations (∼50% of human tumors) or HGSOC lacking a *PTEN* deletion (94% of human tumors). Here, we release a new mouse model of ovarian cancer, driven by p53 mutation and MYC under the Ovgp1 promoter. Evidence of hallmarks of HGSOC and uterine serous cancer are described.

## Methods

### Transgenic OvTrpMyc mouse model creation

Transgenic mice were generated by the Taconic/Cyagen Transgenic Animal Center (Cyagen Biosciences Inc.; Santa Clara, CA). Briefly, *Trp53* (NM_001127233.1) containing an R270H mutation and *Myc* (NM_001177352.1) were knocked into the endogenous Trp53 locus in a reverse orientation using CRISPR-Cas9 mediated gene editing. A construct with 985bp of the murine *Ovgp1* promoter, including its proximal epigenetic regulatory peak [18] (**Figure S1A**), was cloned upstream of the murine *Trp53* coding sequence containing a consensus Kozak sequence and an R270H mutation (CGT to CAT mutation in codon 270) without a stop codon. A P2A self-cleaving peptide sequence precedes the murine *Myc* coding sequence which contains a 3X hemagglutinin (HA) C-terminal tag and is followed by a bovine growth hormone polyadenylation sequence. Homology arms were supplied by bacteria artificial chromosomes RP23-240I24 and RP23-243M15 [19]. The complete transgenic sequence is supplied as Supplementary File 1.gb.

### Mouse husbandry, genotyping, and loss of heterozygosity

All mouse procedures were approved by the Medical University of South Carolina (MUSC) Institutional Animal Care and Use Committee (IACUC), protocol #01305, prior to use. Animals were housed in individual vented cages in rooms with 12:12 light cycle and were fed ad libitum (Purina Pro-Lab 5V75). Founder mice were generated in the C57BL/6Tc background and backcrossed to C57BL/6J for experimentation. Litters were always formed by crossing a transgenic male to a wild-type C57BL/6 mother obtained externally (Jackson Labs, #000664). For genotyping, ∼2 mm sections of mice tails or ear punches were lysed at 55 °C overnight in 350 µL of buffer containing 100 mM Tris-HCl pH 8.8, 5 mM EDTA pH 8.0, 0.2% SDS, 200 mM NaCl and 100 µg/mL proteinase K (VWR 97062-670). After briefly centrifuging to spin down debris, DNA in 300 µL of lysate was precipitated by adding 1 mL of 100% ethanol. DNA was then pelleted by centrifugation at 16,000 g for 30 minutes at 4 °C. The supernatant was discarded, and the pellet was washed with 70% ethanol. After centrifugation at 16,000 g for 20 minutes at 4 °C, the pellet was resuspended in 50 µL of TE (10 mM Tris-HCl, 0.2 mM EDTA pH 7.5). Finally, the tubes were placed in a heat block at 55 °C for 2 h with lids open to evaporate remaining ethanol. Two PCR reactions per mouse were used to distinguish mutant and wild type mice. Both PCR reactions used the same reverse primer (GTGAGATTTCATTGTAGGTGCCAG) and the forward primers were either specific to the knockin construct (CAGTTGAGATGGAGGAGACCTAGG) or the wild type *Trp53* allele (AGCCTGTTGAGCTTCACCCC). For loss of heterozygosity studies, identical genotyping PCR reactions were performed, using genomic DNA extracted from *ex vivo* grown cells via the Invitrogen PureLink Genomic DNA Mini Kit (#K182001).

Matings between male heterozygous OvTrpMyc and female C57BL/6J mice exhibited typical Mendelian inheritance. Of 585 offspring mice, 307 (52%) were male and 278 (48%) were female and 305 (52%) were wild type and 280 (48%) were transgenic. Male mice were 47% transgenic and 53% wild type; female mice were 49% transgenic and 51% wild type. None of these differences were statistically different from the 1:1 ratios predicted by Mendelian segregation.

### Longitudinal study of lethal tumor burden

All mice intended to be studied for end-of-life criteria were carefully monitored for signs of tumor burden starting at 2-months of age. Mice were monitored twice weekly for (1) indications of ascites by swollen abdomen, (2) palpable tumors present externally on the mouse, and (3) weight loss. Euthanasia was performed when mice had a palpable tumor of >1cm, any ascites was observed to increase over two days, gait was impaired, or weight loss surpassed 15% of maximal observed weight. Mice that were found dead for unknown reasons were counted as likely-tumor burdened mice and are included in the Kaplan-Meier curve, but histology was never performed on these mice for quality control reasons. Comparison genotypes were previously published by other labs for p53+/-[20] and Pax8-driven models [17].

### Histopathology

Mice had uterine horns and attached fallopian tubes, ovaries, and fat pads surgically removed. Tissue was fixed overnight in 10% normal buffered formalin (VWR 10790-714), and then transferred to 70% ethanol for 24h. A Shandon Citadel 2000 Processor (Thermo Fisher Scientific) was used to process tissue on a 4-hour cycle with the following reagent cycles: 70% ethanol followed by two changes of 95% ethanol, three changes of 100% ethanol, one cycle in a 1:1 solution of ethanol and xylene, three changes in xylene, and two changes in paraffin. During the embedding process, ovaries with a portion of the fallopian tube were laid down flat, and the remaining sections of the uterus were stood perpendicular for cross sections, averaging 3-4 cross sections of the uterus. After the paraffin hardened, the tissue block was cut in 20µm sections until full sections of ovary and fallopian tubes were visible. Then, 5µm sections were collected onto positively charged slides, air dried at room temperature overnight, and heat fixed at 60 °C for 1 hour before hematoxylin and eosin (H&E) staining or immunohistochemistry. Immunohistochemistry utilized Leica Bond III for automated processing. Antibodies used included rabbit anti-PAX8 antibody (Proteintech 10336-1-AP) at a 1:2000 dilution, rabbit anti-p53 antibody (Leica Biosystems NCL-p53-CM5p) at a 1:5000 dilution, anti-c-MYC antibody (Abcam ab32072) at a 1:500 dilution, and rabbit anti-HA antibody (Cell Signaling 3724) at 1:5000 dilution. Sections were imaged on a BioTek Lionheart FX microscope and images processed using Gen5 and ImageJ software. Mice without visible tumors were necropsied and H&E slides were assessed by a veterinary pathologist for Supplementary Figure 2, and by a gynecologic pathologist for Supplementary Figure 3.

### *Ex vivo* tumor cell culturing conditions

A cohort of mice was used to isolate cells from ovaries, uteruses, or visible tumors. Mice were euthanized and immediately dissected for cell extraction. Ovaries and fallopian tubes were excised from fat pads and uterus. The mid-section of the uterus was used for uterine cell extraction. Visible tumors were excised from surrounding tissue. All tissue was first rinsed once in sterile PBS, and the PBS was then aspirated. Single, ethanol-sterilized razor blades were then used to mince 2mm sections of tissue in a tissue culture hood. Thoroughly minced tissue was then pipetted using a P1000 pipette into a 6-well tissue culture dish, again dispersing cells by repeated pipetting. Cells were initially grown in RPMI 1640 containing glutamine (VWR, #95042-508), supplemented with 20% fetal bovine serum (FBS) (Thermo Fisher, #10437028), 5% penicillin streptomycin solution (Sigma, P4333-100ML), 1% non-essential amino acids (Sigma, #TMS-001-C), and 1% sodium pyruvate solution (Sigma, #S8636-100ML). After 48h, media was aspirated and replaced with similar media containing only 2% penicillin streptomycin solution. Media was then replaced every 3-4 days until the dish became 33-67% confluent. Cells were then trypsinized with Trypsin-EDTA (Sigma, #T3924-100ML) for 3-10 minutes until non-adherent, combined 1:1 with complete RPMI, and cell suspension centrifuged at 1,000g for 2 min. Cells were transferred to a new 6-well dish and allowed to become confluent. Once confluency was reached, cells were transferred to larger plates for culturing and freezing aliquots in 10% DMSO. Once cell lines were established by growing in culture for a minimum of two weeks containing two trypsinization events, cells were grown as cell lines in complete RPMI. Complete RPMI consisted of 500ml RPMI 1640 containing glutamine, 5ml of sodium pyruvate solution, 5ml of penicillin streptomycin solution, and 50ml FBS. Cells were passaged within a day of surpassing 80% confluence by 1:10 dilution.

### Transcriptome sequencing and comparative analysis

RNA was extracted from cells using a Qiagen RNeasy Miniprep kit (#R1054). RNA quality was assessed using an Agilent 4200 TapeStation and RINe values ranged from 8.6-10. Polyadenylated RNA was captured from 500ng of total RNA per sample, and libraries were prepared using the NEBNext Poly(A) mRNA Magnetic Isolation Module (#7490L) and NEBNext Ultra II Directional RNA Library Prep Kit for Illumina (NEBNext, #7760L). Paired-end sequencing was done at the Vanderbilt VANTAGE core laboratory (Vanderbilt University) to a depth of 25 million reads per library using an Illumina NovaSeq 6000. Raw fastq.gz data were uploaded into Galaxy [21]. GRCm38.86 (mm10) gtf.gz and toplevel.fa.gz reference files were obtained from Ensembl [22]. RNA STAR [23] was used to align paired end 150bp reads to mm10, followed by PCR duplicate removal by SAMtools RmDup [24]. Mapped reads were attributed to transcripts using featureCounts [25] and quantified for gene and transcript expression fragments per kilobase of transcript per million reads mapped (FPKM) using Cufflinks [26]. Expression of CGT>CAT mutant p53 was manually verified in read data using mapped read visualization in the UCSC Genome Browser mm10 database around region chr11:69,589,594-69,589,622. One of eleven samples from an *ex vivo* cell line did not retain expression of the mutated copy of p53 (F361 tumor cells), despite having mapped RNA reads to this codon. FPKMs per sample were combined into transcripts per million (TPM) reads tables using R to enable normalized sample-to-sample comparisons [27].

For human cancer cell lines to compare with, human Cancer Cell Line Encyclopedia [28] FPKM expression data were used and converted to TPMs. To enable cross-species comparison, reciprocal orthologs data were utilized. Mousemine.org [29] and Humanmine.org [30] were used for ortholog identification and the intersect of genes were used as reciprocal orthologous genes, resulting in 13,229 genes expressed and common orthologs between the two datasets. Human cell lines compared to mouse cancer cell lines included all lines annotated as from cervix, endometrium, and ovary (N= 80 cell lines). Transcriptomes were pairwise compared using a Euclidean distance metric of all expressed genes. Nearest neighbor human cell lines comprise the cell lines shown. Gene expression in this figure was further normalized for visual comparison through mean transcription normalization, including the additional cell lines HELA, KURAMOCHI, SIHA, A2780, CAOV4, MDA-MB-231, MDA-MB-453, and KHYG cell lines, chosen as representative cell lines which have mutational characteristics as meaningful to compare with the mouse cell lines (see Results). Shown signature genes were the top 25 genes with minimal expression differences within the mouse cell lines and nearest-neighbor human cell line and maximal differences with non-nearest neighbor cell lines. DAVID [31] was used to generate the pathway annotations shown in the transcriptional signature.

### Copy number alteration analysis

Raw counts from featureCounts were input into InferCNV to assess copy-number alterations. Settings for InferCNV included i3 copy number variant (CNV) calling, cutoff value of 1, Hidden Markov Model (HMM) transition probability of 1e-02, and a window length of 300. Samples F326 splenic tumor, F326 ROV, and F361 tumor were used as control inputs due to lower levels of initial CNV calls in these samples.

### Syngeneic intraperitoneal modeling

Cells, lentivirally labeled with GFP, were grown to 5-10 million per mouse and injected in 200µl iced PBS via intraperitoneal injection. Recipient mice were 3-month-old female OvTrpMyc mice. Mice were monitored for euthanasia criteria matching the longitudinal study. Tumors, uterine horns, and attached fallopian tubes, ovaries, and fat pads were then immediately fixed and processed for immunohistochemistry.

### Data availability

Next generation sequencing raw and processed data is available via GEO accession GSE238158.

## Results

### Generation of a human-mimetic genetic model of HGSOC and serous endometrial cancer

Previous models of epithelial ovarian cancer have been used to seek a better understanding of the disease using known mutations. Ovarian cancer most commonly presents as high-grade serous ovarian carcinoma (HGSOC), which contains p53 mutation in at least 94% of cases [32, 33], sporadic *BRCA1* or *BRCA2* mutations in <15% of cases. Germline *BRCA1* or *BRCA2* mutations yield ovarian cancer, most often HGSOC, with a lifetime unmitigated risk of 30-70% [34]. Other mutations are present in 15% or less of patient primary tumors, including *APC, MUC16, ARID1A*, *RB1*, *NF1*, *CSMD3*, *CDK12*, *FAT3*, and *GABRA6* [32, 35]. However, HGSOC exhibits unusually high levels of copy-number alterations (CNAs) and aneuploidy, affecting two-thirds of all genes. Low grade serous carcinomas are driven by *BRAF* and *KRAS* and represent <5% of ovarian tumors [36]. Mucinous tumors are driven by *KRAS* or *ERBB2* amplification, representing about 3% of ovarian tumors. Endometrioid and clear cell carcinoma each represent about 10% of ovarian tumors and have high rates of *ARID1A* mutations (30 and 50%, respectively) and *PTEN* alterations including deletions and mutations. Clear cell carcinomas mutate *PIK3CA* in 33% of tumors. Mouse models of ovarian cancer have utilized these known mutations, which are reviewed by Howell in 2014 [37] and Zakarya *et al* in 2020 [38]. A comparison of these mouse driver genes to known human genetic driver frequencies is shown in **Figure 1A**. Notably, uterine serous carcinoma, a type II endometrial cancer, exhibits very similar mutation patterns to HGSOC.

**Figure 1.**
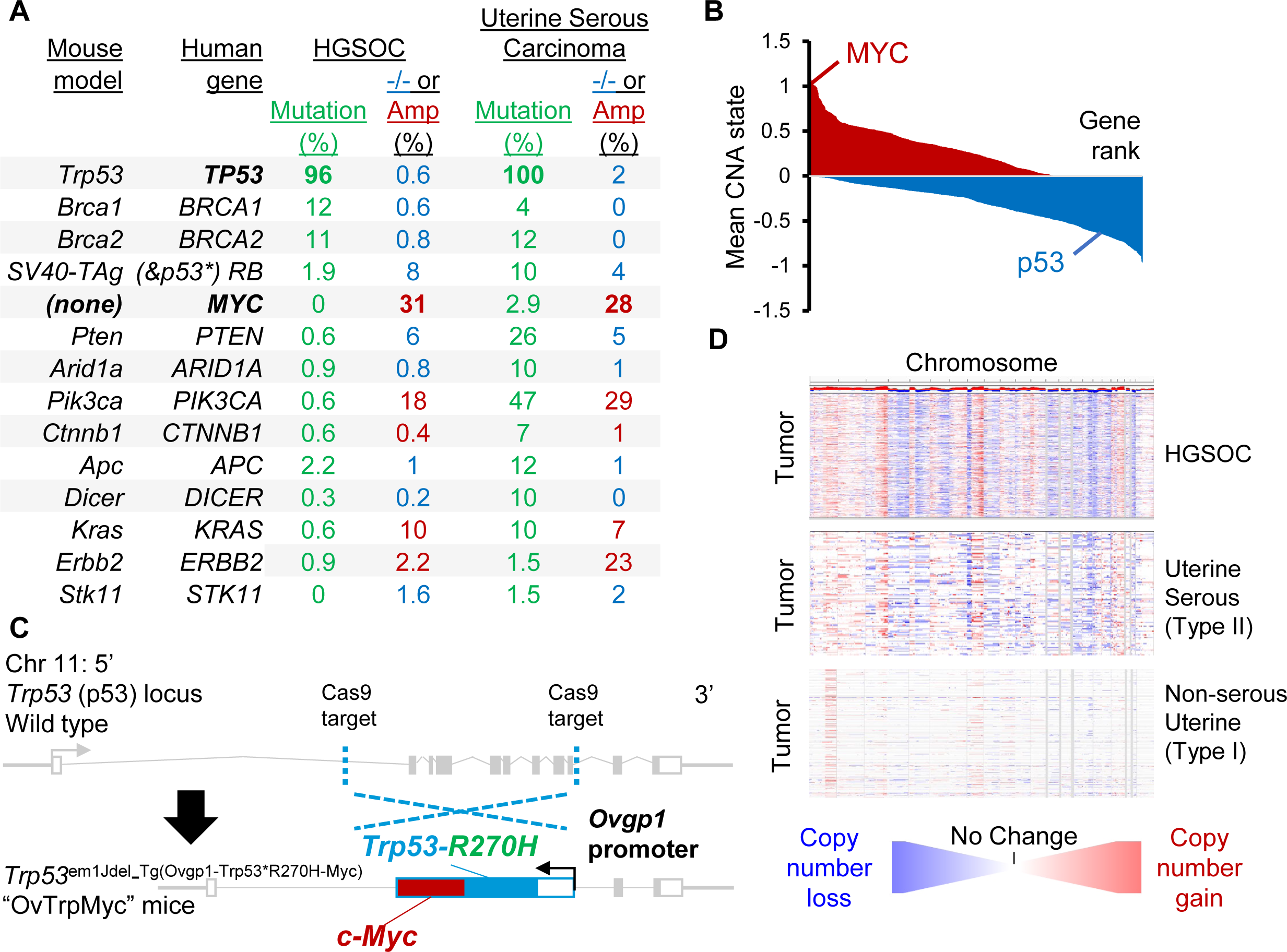
Rationale and design of transgenic Ovgp1-Trp53-R270H-Myc mouse model. **(A)** Comparison of currently available HGSOC mouse models genetics to genetic alterations in human HGSOC and serous endometrial cancers. -/- refers to homozygous deletion (blue, tumor suppressors) while Amp refers to copy number amplification of at least two extra copies (red, oncogenes). **(B)** Average copy-number state using TCGA HGSOC data at the gene level. Genes are plotted by ranking, with *MYC* and *TP53* highlighted. **(C)** Design of transgenic OvTrpMyc mouse model. The endogenous Trp53 allele was targeted using a spCas9 and homology donor repair template strategy in C57BL6/Tc background. Transgene genotyped pups were backcrossed with C57BL/6J mice and heterozygous mice used in all studies. The *Ovgp1*-driven transgene includes a dominant negative murine p53-R270H mutant sequence and a murine *c-Myc* sequence separated by a P2A self-cleaving peptide. **(D)** Comparison of copy-number alterations across the genome for TCGA studied HGSOC and serous UCEC patient tumors. Red indicates copy number gain while blue indicates copy number loss.

We sought to create a new genetic model, Trp53^em1Jdel_Tg(Ovgp1-Trp53*R270H-Myc)^, nicknamed OvTrpMyc, which accurately depicts *BRCA* and *PTEN* normal HGSOC. We and others have found that the most commonly and excessively amplified region of the HGSOC genome is 8q24 [14, 32] (**Figure 1B**). Expression of a strong oncogene located in 8q24, *c-Myc,* was coupled with the expression of mutant p53. MYC is a basic helix-loop-helix leucine zipper transcription factor that regulates cell stemness, metabolism, apoptosis, and cell cycle progression [39]. Ovarian cancers require MYC for proliferation [40]. The most common mutation of p53 in HGSOC [32] is a dominant negative, gain-of-function [41, 42] amino acid mutation R273H, which maps to R270H in mice. We engineered mice via CRISPR-Cas9 targeting to insert a transgene driven by *Ovgp1* into the native *Trp53* (p53) locus of chromosome 11 (**Figure 1C**). *Ovgp1* was previously demonstrated to express specifically in non-ciliated fallopian fimbriae cells; not on the uterine epithelium and not on the ovarian surface epithelium [11, 12]. We confirmed FTE-specific transgene staining patterns within the OvTrpMyc model (**Figure S1B**). The transgene was inserted at the p53 locus to allow for CNAs to evolve including on the p53 resident chromosome, as CNAs are characteristic of HGSOC. All transgenic mice in this study were genotyped as heterozygous for the transgene. HGSOC CNAs are similar to uterine serous carcinoma, but not type I uterine cancer (**Figure 1D**).

### Middle age disease presentation

The median age of tumor-burden induced euthanasia for the OvTrpMyc mice was 15.1 months (**Figure 2A**), or, given the limitations of human-mouse comparisons, about 40-60 years of human age (based on life expectancy of 80 years old, the 896 days female C567BL/6 lifespan [43], and that peri-menopause in mice occurs around 9-months of age [44]). *<u>Pten</u>*-null mouse models drive disease in young mice and previously published survival data are shown for qualitative comparison [17]. Since OvTrpMyc mice are effectively *Trp53+/-* due to targeting the transgene at the native p53 locus, previously published data for *p53+/-* mice are also displayed [20].

**Figure 2.**
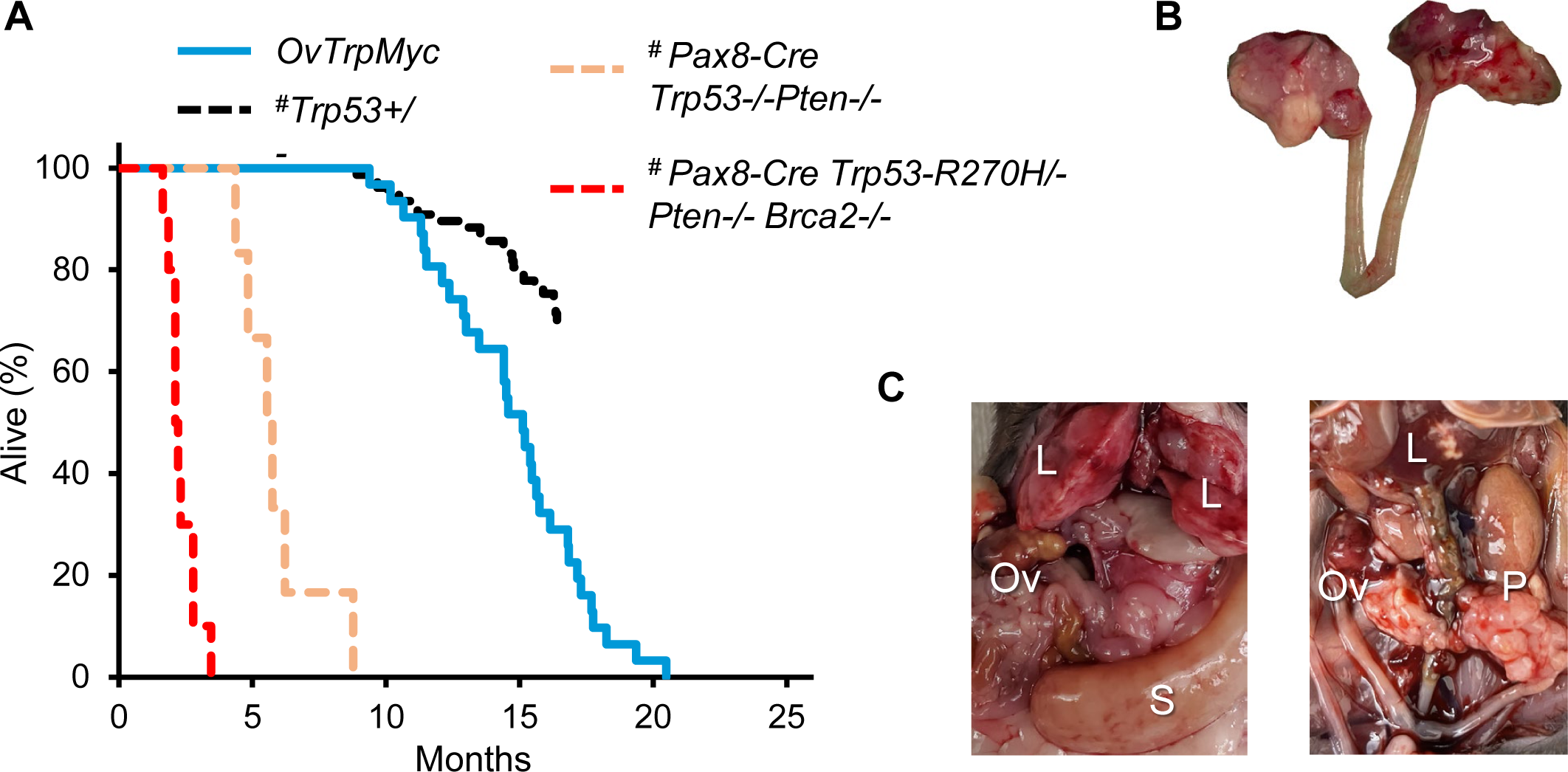
Middle age presentation of ovarian and peritoneal tumors. **(A)** Lifespan data from mice euthanized due to tumor burden criteria. # indicates the Pax8 and p53+/-model survival curve data were assessed from previous publications (see Methods) and are displayed for qualitative comparison. OvTrpMyc data originate from this study. **(B)** example of a dissected reproductive tract from a mouse with obvious macroscopic ovarian tumors. **(C)** examples of widespread intra-abdominal disease, including liver invasion (L), peritoneal tumors (P), splenic tumor (S), and ovarian tumors (Ov).

These mice were designed with limited oncogenic stimuli with the intent of long latency, consistent with human disease. However, OvTrpMyc mice are effectively *Trp53^+/-^* in all cells. Heterozygous *Trp53^+/-^*mice develop lymphomas, osteosarcomas, fibrosarcomas, hemangiosarcomas, lung adenocarcinomas, and other tumors, which form lethal tumors in 25% of mice requiring euthanasia as early as 16.5 months [20]. Complete knockout *Trp53^-/-^* mice develop primarily lymphomas and have a half-life of 4.5 months. We observed our OvTrpMyc mouse model to have a survival curve intermediate between these genotypes (**Figure 2A**).

### Pathology of lethal ovarian and uterine cancers

Gross examination of mice meeting euthanasia criteria showed HGSOC and other cancer phenotypes. For HGSOC, tumors of the ovary were occasionally observed (**Figure 2B**), but large tumors were rare. Widespread intra-abdominal disease was occasionally observed (**Figure 2C**), with tumors throughout the mesentery as well as the peritoneal wall. Tumors involved with muscle external to the abdomen were observed, typically with invasion spreading from the peritoneal wall or from the ovary.

Previous, carefully controlled studies demonstrated *Ovgp1* expresses on non-ciliated epithelial cells within the FTE, not the OSE [11, 12]. To investigate OvTrpMyc transgene expression at end stage disease, the gynecologic tract (including uteri, fallopian tubes, ovary, and ovarian fat pad) was dissected from mice meeting euthanasia criteria at 10-16 months of age and processed for molecular pathology. Tissue was formalin fixed and embedded in paraffin prior to sectioning and staining for p53 and MYC. Tumors, if observed, were additionally stained with paired-box transcription factor PAX8, a biomarker used to help define gynecologic tumors in humans [45–47], but is also expressed in embryogenesis of the thyroid, renal, and upper urinary tracts. Wild-type mice did not exhibit clusters of p53 or MYC positivity on the FTE, although scattered single cells were positive (**Figure 3A**). In single sections, some FTEs from OvTrpMyc mice were also negative, like wild-type, perhaps since the entire fallopian tube was not examined due to limited material availability (**Figure S2**). However, 60% of mice requiring euthanasia exhibited positive FTE staining with p53 and MYC in the single section evaluated. This contrasts with young mice: in three out of three 3-month old OvTrpMyc mice, none showed positive staining, despite clear transgene expression by 10-weeks of age (**Figure S1b**). Morphology and cellular distribution of p53 and MYC varied between mice. Mouse F135 exhibited nuclear p53 and nuclear MYC within the FTE, indicative of HGSOC, with surrounding cells exhibiting a mix of nuclear and cytoplasmic staining (**Figure 3B**). This mouse had a large retroperitoneal tumor extending from the ovary to the spinal cord. The tumor stained positive for PAX8 in addition to p53 and MYC (**Figure 3C**), as did a small mass situated in the ovarian fat pad (**Figure 3D**), consistent with known HGSOC homing to adipocytes [48]. Other mice exhibited FTE staining with primarily cytoplasmic expression surrounded by vacuoles (**Figure 3E**).

**Figure 3.**
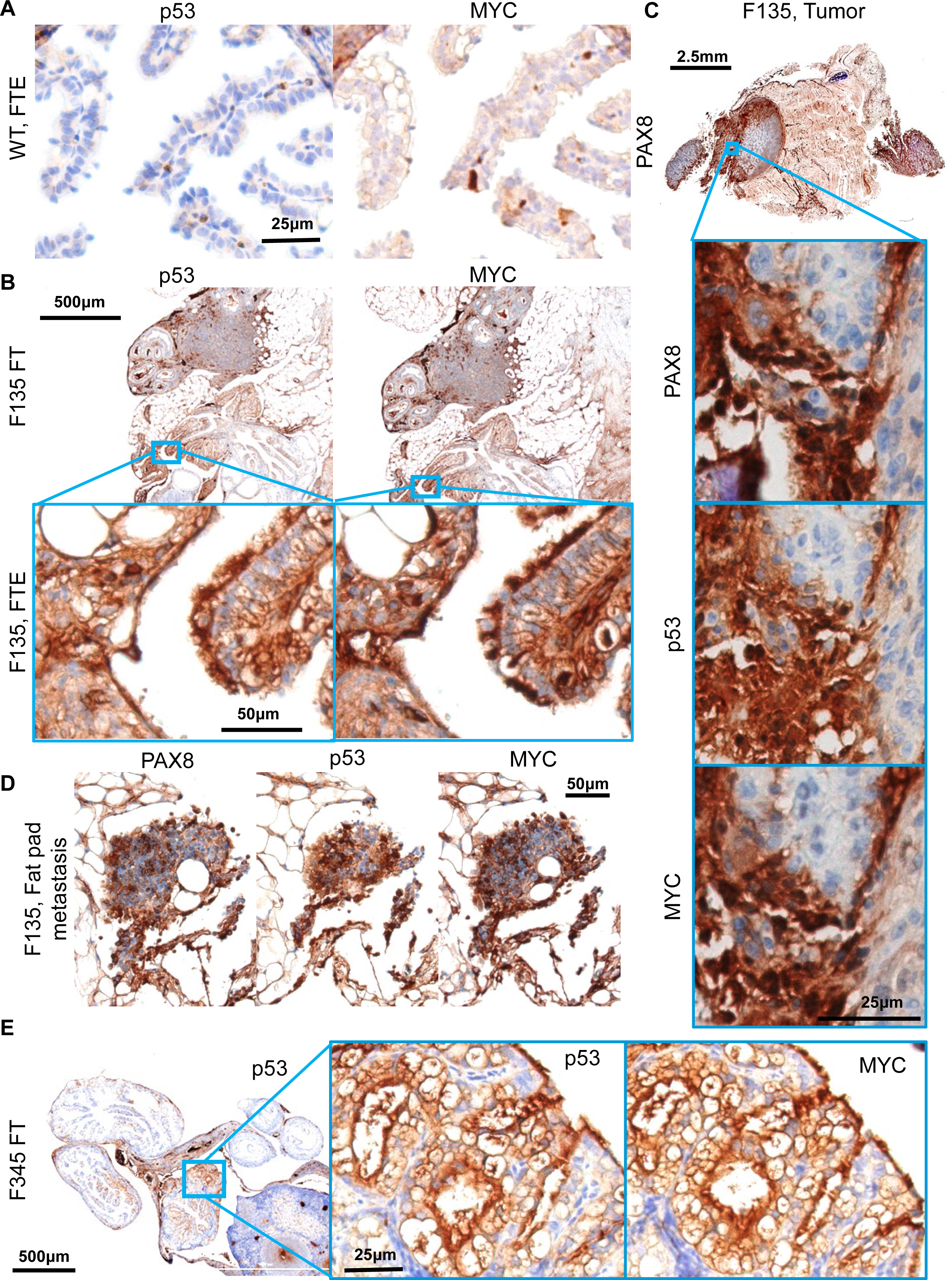
Molecular histology of fallopian fimbriae. **(A)** A wild-type mouse at 12mo of age shows little p53 or MYC staining in fallopian tube epithelium (FTE). **(B)** OvTrpMyc fallopian tube (FT) at 15mo of age, F135, exhibiting p53 and MYC staining within FTE. **(C)** Intra-abdominal tumor dissected from an OvTrpMyc mouse with positive PAX8, p53, and MYC staining. **(D)** An ovarian fat pad metastatic nodule. **(E)** OvTrpMyc mouse at 12mo of age, with vacuolated fallopian fimbriae positive for p53 and MYC.

OSE staining of p53 and MYC was observed in 50% of OvTrpMyc mice studied. In OvTrpMyc mouse F278, nuclear positivity of p53 and MYC was observed in cell clusters on the OSE (**Figure 4A**). Mouse F278 exhibited minimal cytoplasmic positivity in the FTE and uterine luminal epithelium. The intra-abdominal tumor found in mouse F278 displayed PAX8 staining in addition to p53 and MYC (**Figure 4B**). Other transgenic mice, similar to wild-type, were not observed to have p53 or MYC positivity on the OSE. Like wild-type mice, other ovarian structures stained positive (**Figure 4C,D**). The positive OSE observations may be intraepithelial metastasis events, given the lack of *Ovgp1* expression in the OSE.

**Figure 4.**
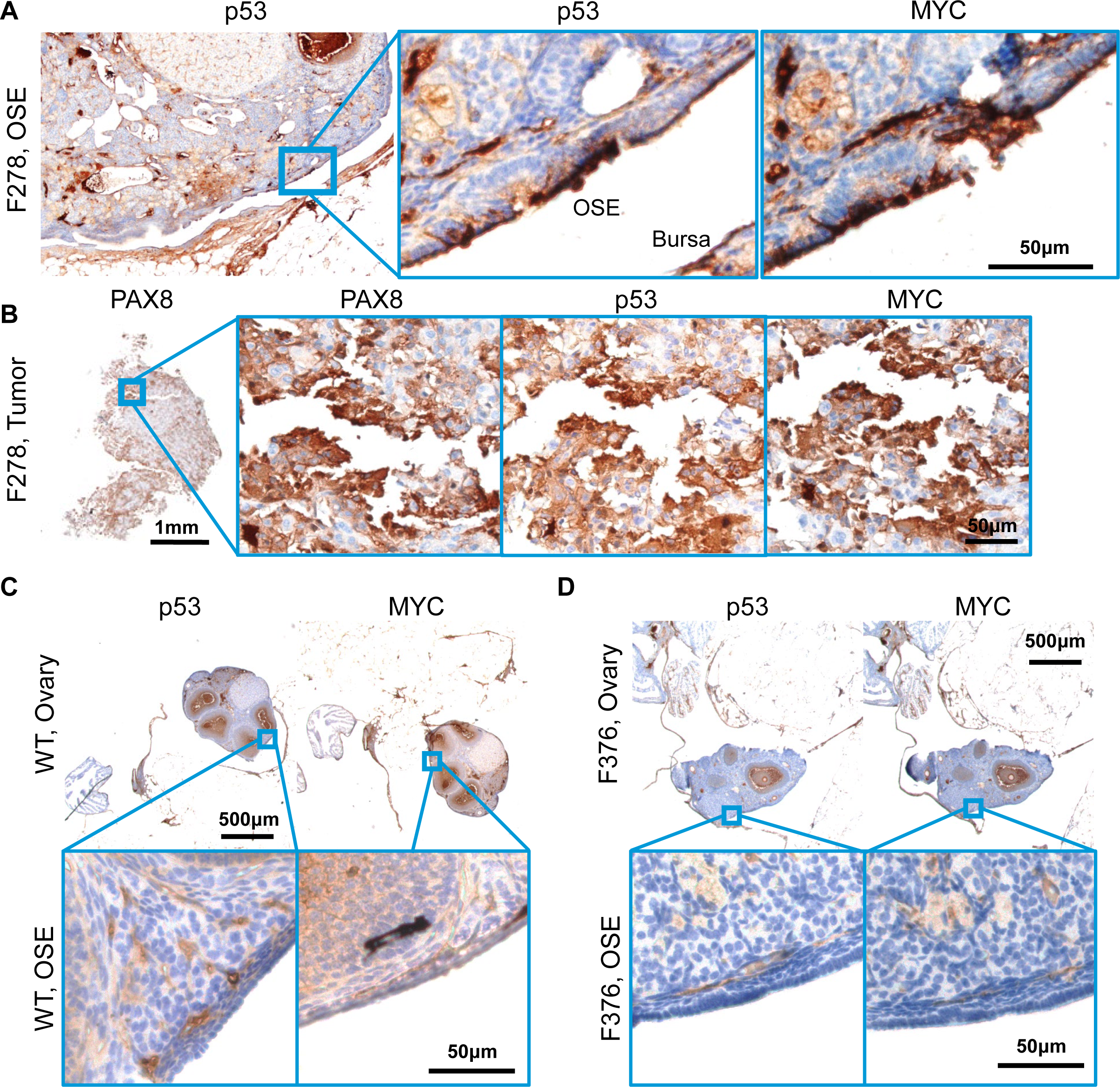
Molecular histology of ovarian surface epithelium. **(A)** An OvTrpMyc at 16mo of age with clear ovarian surface epithelium (OSE) staining of p53 and MYC. **(B)** The intra-abdominal tumor extracted from the mouse shown in (a) exhibited PAX8, p53, and MYC staining. **(C)** A wild-type mouse at 12mo of age is negative for p53 or MYC staining in OSE. **(D)** Example of an OvTrpMyc mouse with negative OSE staining of p53 and MYC, at 12mo of age.

While the OvTrpMyc mice were originally designed to mimic HGSOC, strong staining of p53 and MYC on the uterine luminal epithelium was noted in 60% of OvTrpMyc mice. While adjacent tissue exhibited p53 and MYC staining in normal, wild-type mice, uterine epithelium was universally negative for staining (**Figure 5A**). In OvTrpMyc mice, positive epithelial clusters were observed, flanked by scattered cells evidently spreading into adjacent epithelium, which was not observed in wild-type mice (**Figure 5A,B**). This occurred in the absence of OSE (**Figure 4D**) or FTE staining (**Figure S2**) in the single section from OvTrpMyc mouse F376, but coincided with macroscopic tumor staining (**Figure 5C**). However, uterine lumen epithelial staining coincided with predominantly cytoplasmic p53 and MYC staining found in the OSE in one of the youngest animals euthanized, at 11 months (**Figure 5D,E**). Ovarian fat pad metastatic cells were found and included p53, MYC, and PAX8 staining (**Figure 5F**).

**Figure 5.**
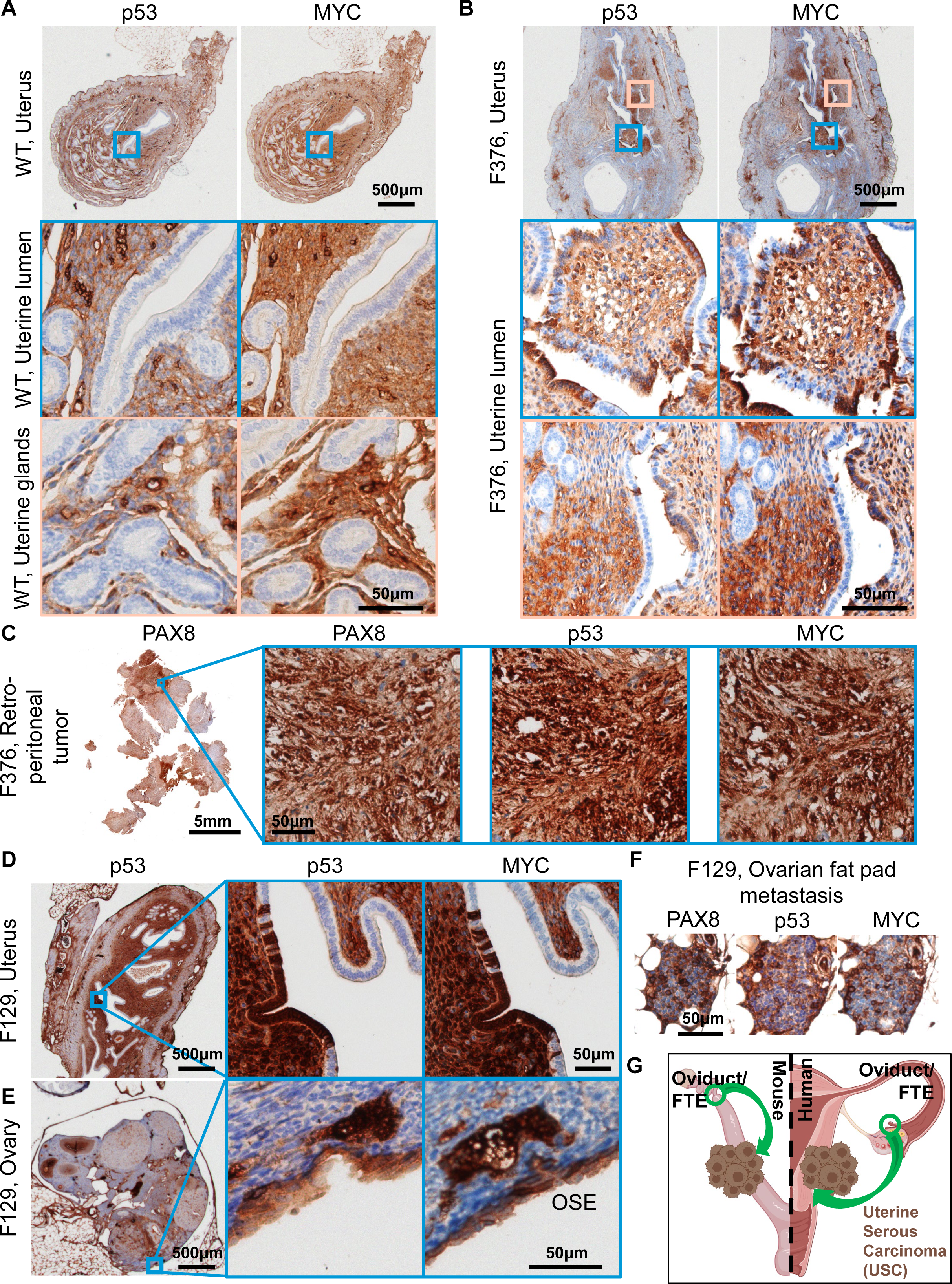
Uterine luminal epithelium staining of p53 and MYC. **(A)** A wild-type mouse at 12mo of age shows negative p53 and MYC staining in uterine epithelium and positive staining in adjacent tissue. **(B)** Example of a 12mo old OvTrpMyc mouse with cytoplasmic p53 and whole-cell MYC staining along uterine luminal epithelium. **(C)** Large retroperitoneal tumor, behind kidney, stained for p53, MYC, and PAX8. **(D)** An example of uterine luminal epithelium expression in OvTrpMyc mouse F129, with evidence of similar whole-cell staining of p53 and MYC on the **(E)** OSE and **(F)** ovarian fat pad metastasis. Mouse F129 was 11mo of age. **(G)** Proposed model for observed dissemination of cells from FTE to uterine luminal epithelium as an origin of uterine serous carcinoma

### Observation of non-gynecologic cancers

Macroscopic tumors were not always immediately observed upon dissection, despite a mouse reaching euthanasia criteria. Three such mice were subjected to full necropsies, revealing brain tumors and lung tumors (**Supplementary** Figure 3). Tumors resembling sarcomas or carcinosarcomas were also identified from macroscopic neoplasms (**Supplementary** Figure 4). It is unclear if these are metastatic events, a form of uterine carcinosarcoma, accelerated background strain tumor formation due to disruption of *Trp53*, or another event.

### Aneuploidy and copy-number alterations within OvTrpMyc tumor cells

The primary rationale of integrating the OvTrpMyc transgene into the endogenous p53 locus was to allow for loss of heterozygosity (LOH) of chromosome 11 (chromosome 17, in humans) during spontaneous tumor development. To test if LOH occurred in tumor cells from these mice, eight mice meeting euthanasia criteria were dissected for small portions of the uterus, ovaries, fallopian tubes, and any macroscopic tumor presentation. These dissections were minced, washed, and allowed to grow *ex vivo* in tissue culture media (see Methods). Once a cell line was established with regular passaging, these cells were (a) genotyped for LOH and (b) sequenced by bulk RNA-seq for mRNA transcriptome analysis. Seven mice yielded a cell line from at least one of these origin sites. LOH was present in the *ex vivo* OvTrpMyc cell lines generated (**Figure 6A**; mutant PCR bands were seen, but no wild-type bands were seen other than in controls). These results provide direct evidence that the wild-type allele of p53 was lost in these tumor cells. Each tumor cell line was shown to have ample chromosome alterations (**Figure 6B**), consistent with patterns in human disease. While synteny is partial for most chromosomes, loss of the p53-coding chromosome (11 in mice, 17 in humans) and gain of the mouse region syntenic to human amplicon 8q24, on murine 15qD1, was often found.

**Figure 6:**
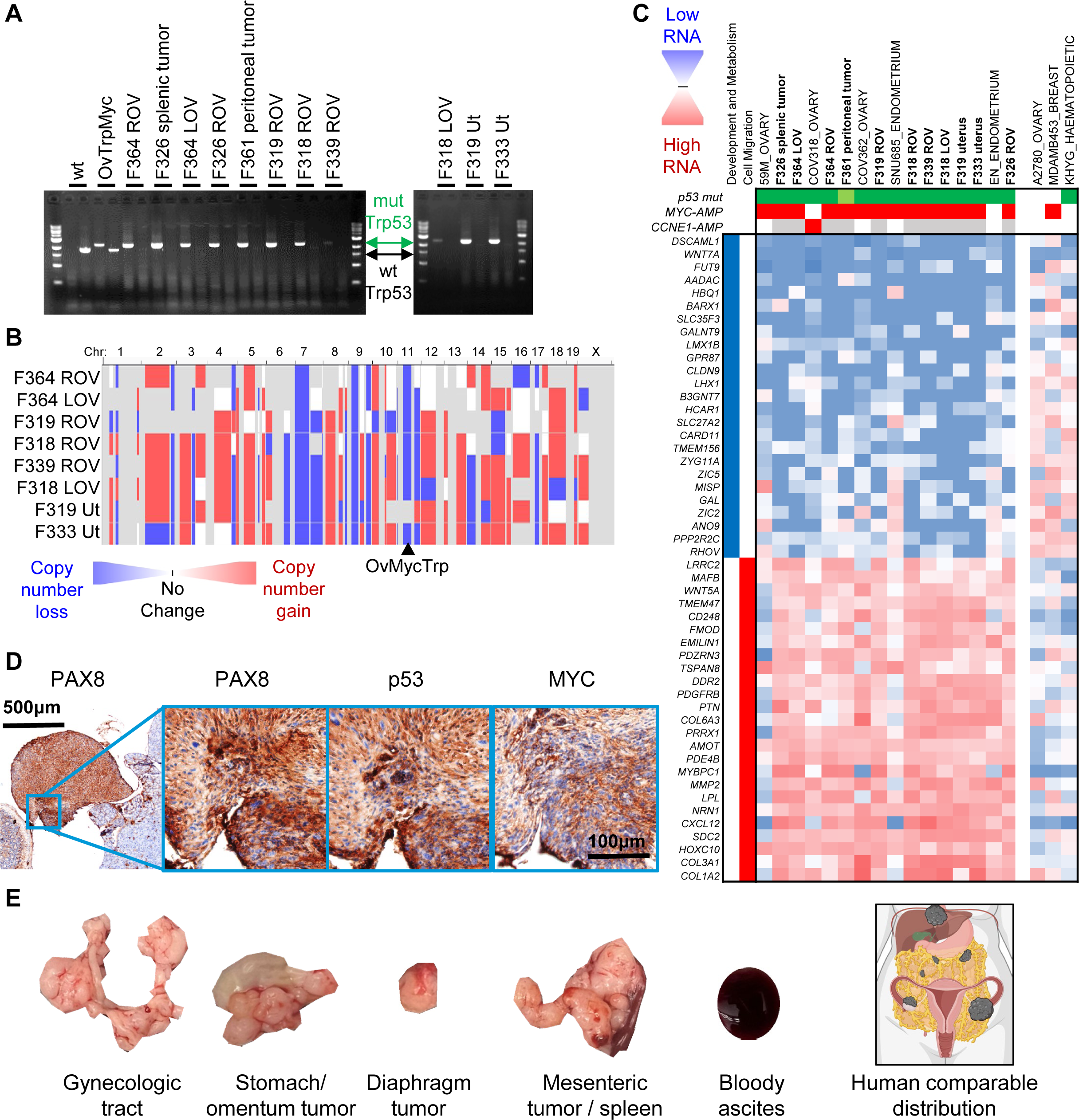
Copy number alterations, transcriptomics, and tumorigenicity of extracted OvTrpMyc tumor cells. Label key: LOV= left ovary, ROV= right ovary, Ut=uterus **(A)** Loss of heterozygosity was observed by PCR of the Trp53 transgene (mut) in isolated cells. **(B)** Whole genome copy number alteration calls of tumor cells. Blue indicates copy-number loss and red indicates copy-number gain. **(C)** Transcriptomic profiling of OvTrpMyc cancer cells compared to human gynecologic cancer within the Human Cancer Cell Line Encyclopedia. Header genes refer to mutation in p53 (green), MYC amplification (red), or CCNE1 amplification (red), shaded white if absent, or gray if unknown. Nearest neighbor human cell lines comprise the cell lines shown. A2780, MDA-MB-453, and KHYG cell lines are shown as representative cell lines which have driver gene characteristics comparable with the mouse cell lines, but6 clearly different transcriptional profiles. **(D)** 8x10 F339 ROV OvTrpMyc cell line cells were injected IP into a young OvTrpMyc female recipient mouse. The mouse lost weight, meeting euthanasia criteria after 54 days. Histochemical staining of a dissected tumor attached to the intestines is shown. **(E)** Distribution of6 tumors found in F318 LOV IP syngeneic injected mice (5x10 cells, 132 days), relative to a human cartoon example of HGSOC metastatic spread.

### OvTrpMyc tumor cells transcriptionally pair with HGSOC and uterine serous cancer

Transcriptomes from OvTrpMyc tumor cells were next analyzed to characterize which human gynecologic cancers they best mimic. Since cell lines have clear differences from endogenous disease [49], transcriptomes we compared between mouse and human cell lines. Clustering of transcriptomes revealed that tumor cells from OvTrpMyc mice most closely mimicked well characterized HGSOC and p53-mutant endometrial cancer cell lines, not cervical nor other subtypes of ovarian cancer (**Figure 6C**). A singular exception in cell identity clustering was the EN cell line, which in the Cancer Cell Line Encyclopedia (CCLE) dataset, utilized for transcriptomes in Figure 6C, was not annotated as p53 mutant nor *MYC* or *CCNE1* amplified. However, the founders of the EN line note a p53-R273H mutation [50], and the transcriptomic data from the CCLE [51] also show *MYC*, *CCNE1*, and p53 expression as comparable to HGSOC lines. Cell lines with features that are only a portion of the transgene are shown for comparison. A2780 is a non-HGSOC ovarian cancer cell line, but has neither a p53 mutation nor a MYC amplification. Triple negative breast cancers have many similar characteristics from a genetic standpoint to HGSOC including frequent BRCA mutation, p53 mutation, and high aneuploidy, but a representative cell line MDA-MB-453 displayed a clearly different transcriptomic profile. Lastly, the leukemia cell line KHYG, which has a p53 mutation, also displayed a clearly different transcriptomic profile. OvTrpMyc clusters revealed a downregulation of developmental and metabolic genes and an upregulation of cell migration genes. These transcriptome-wide data provide molecular evidence that complements histopathology to confirm HGSOC and serous endometrial cancers arise in the OvTrpMyc GEMM.

Cell lines were amplified and injected intraperitoneally into recipient young female OvTrpMyc mice. Mice were monitored for disease meeting the same euthanasia criteria as the spontaneous disease model. Mice required euthanasia due to ascites or distorted abdomen at 40 days (F319 uterus), 54 days (F339 ROV), 65 days (F326 ROV), and 132 days (F318 LOV). Upon euthanasia, gross pathology observations were noted, the gynecologic tract dissected, and tissues were processed for histopathological analysis. Histopathological examination of intra-abdominal tumors showed evidence of HGSOC, including nuclear atypia and pleomorphism, cellular budding, PAX8 staining, nuclear p53 staining, and nuclear MYC staining (**Figure 6D**). These F339 ROV and F318 LOV cells are evidently useful for syngeneic injections for controlled experimentation regarding HGSOC tumor formation and development. The F319 uterus cell line exhibited rapid ascites and may be considered for uterine serous carcinoma syngeneic experiments.

## Discussion

Our study demonstrates that MYC expression and a *p53-R270H* mutation drives HGSOC in a mouse model of Ovgp1-driven fallopian tube epithelium transgene expression. No *BRCA* or homologous repair gene was altered in the GEMM design. The single transgene is sufficient for experimentation without extensive breeding protocols. Latency is long, 15.1 months on average, allowing for the development of HGSOC characteristics such as peritoneal metastasis and extensive copy-number alterations. Intraepithelial metastases to the ovarian surface epithelium and uterine luminal epithelium were observed. Cells grown *ex vivo* from the gynecologic tract from transgenic mice formed tumors following intraperitoneal injections. This mouse model may be used to study aspects of spontaneous cancer development and for syngeneic injection experimentation.

At morbidity, 100% of studied mice exhibited evidence of p53 and MYC positivity along epithelial clusters in at least one site within the FTE, OSE, and uterine luminal epithelium. 40% exhibited positivity at multiple sites. Yet, the largest tumor observed in these mice tended to be metastases, not the ovary itself. We speculate that this may be due to high metastatic potential of these cells which have both *p53* mutation and MYC overexpression, consistent with the finding that syngeneic mouse injections yielded lethal tumor burden. Given that *Myc* is a Yamanaka factor enabling de-differentiation and re-differentiation into many tissue sites [52], it may be unsurprising that metastases predominate in this GEMM.

The OvTrpMyc mice have a longer latency than other models with strong drivers. SV40 T-antigen MISIIR mice develop ovarian tumors with complete penetrance, usually within 4-6 months of age [53]. With *Ovgp1*, SV40 driven mice develop histological lesions in the epithelium of the fallopian tube as early as 6 weeks of age [54]. Mice with *Pax8-Cre* driven *PTEN* knockouts were lethal at 2.2-5.7 months, from disease originating in fallopian STICs [17]. Similar to *Brca1-/-p53-/-Rb-/-Nrf1+/-* mice [12], the OvTrpMyc mice often require a year or more to develop observable disease. We predicted latency is required to accumulate aneuploidy, which we previously observed was somewhat rare in SV40 T-antigen driven disease [14]. Aneuploidy is characteristic of HGSOC and uterine serous cancer [32, 55, 56]. A handful of aneuploid events were observed previously in *Brca2-/-Pten-/-p53^R270H/-^* driven tumors [17]. Here, OvTrpMyc mice clearly utilize aneuploid mechanisms for tumor development processes, with strong positive selection for the mutant allele of p53 on chromosome 11. CNAs were ubiquitous in cell lines derived from OvTrpMyc.

Fallopian fimbriae as a likely origin site of ovarian cancer was first suggested from histopathology of prophylactic oophorectomy from women with germline BRCA mutations [57]. GEMMs have now provided ample support for epithelium within the fallopian fimbriae as a site with characteristics of disease initiation [37, 38]. However, *Ovgp1* FTE mouse models all included a *Brca1+/-* or *Brca1-/-* genotype along with other driver genes. There remains clinical debate as to whether salpingectomies should be performed in patients of average risk, particularly opportunistic procedures during tubal ligation for contraception [10, 58]. The OvTrpMyc model presented here is the first HGSOC GEMM showing evidence of precursor lesions in the fallopian fimbriae epithelium using the *Ovgp1* promoter without *Brca* deletions. This is supportive of the thesis that patients without germline *BRCA* mutation may nonetheless benefit from salpingectomies to reduce ovarian cancer risk.

Limitations of this model are varied. Some of the non-gynecologic tumors found in late-life euthanized mice were analogous to those found in *Trp53+/-* mice. While we also found that gynecologic cancer cells could be isolated in 88% of cases of a dissected mouse at end-stage disease, metastatic modeling of HGSOC in this spontaneous model is difficult. PAX8 and other biomarker staining can aid in interpretation of site of origin. Mice envelop the ovary with a layer of tissue forming the bursa, which humans do not have, and this may affect metastatic properties. The bursa is additionally known to develop leiomyosarcomas in response to intrabursal viral inactivation of *Trp53* and *Brca1* [59].

The unexpected, yet exciting, finding that hallmarks of uterine serous carcinomas, such as cells resembling endometrial intraepithelial carcinoma, are found in the OvTrpMyc model provides further support that uterine serous cancer is similar genetically and biologically to HGSOC. Like HGSOC, the most common p53 alteration in uterine serous carcinomas is a mutation in R273 (murine R270). For CNAs, 8q24, containing *MYC*, is the second most common amplification just behind the unstable *MECOM* locus near *PIK3CA* [55]. Previous serous endometrial GEMMs required loss of *PTEN* in a p53 null background or a mutation in a telomere shelterin complex gene to yield disease within the lifetime of a mouse [60, 61]. Uterine serous carcinoma is aggressive and treatment resistant like HGSOC; it only accounts for 10% of endometrial cancer cases, yet it is responsible for 39% of the deaths [62]. Retrospective studies have revealed a link between tubal ligation, which may prevent intraepithelial metastasis from the fallopian tube, and reduced rates of endometrial cancer [63]. In a carefully controlled cohort study of tubal ligation, the hazard ratio for Type I endometrioid uterine cancer was 0.78, but the hazard ratio was strikingly 0.25 for Type II, typically p53-mutant uterine serous carcinoma, indicating surprisingly low incidence of Type II tumors following tubal ligation [64]. Fallopian STICs and other fallopian epithelium irregularities were observed in uterine serous carcinoma patients, and included the same clonal p53 mutation [65, 66]. Taken together, these human retrospective studies, the current OvTrpMyc GEMM study, and human genetic data [67, 68] are all consistent with the emerging hypothesis that uterine serous carcinoma can originate from the same site as HGSOC: the FTE.

## Acknowledgements

Assistance in the preparation and specimen staining were provided by the Histology and Immunohistochemistry Laboratory, Dept. of Pathology and Laboratory Medicine, Medical University of South Carolina. We thank Audreanna Miserendino and Sydney Oesch for helpful comments on manuscript preparation. We thank all private donors, particularly Matt Prisby for organizing the Sheryl Prisby gynecologic oncology fund, for their individually perhaps small yet cumulatively enormous contributions to advancing cancer research and health. We thank the Division of Laboratory Animal Resources staff, Jackson laboratories staff, and Cyagen/Taconic staff for technical aspects of mouse care and transport. Funding sources. Supported in part by a grant from the Rivkin Center for Ovarian Cancer. Research reported in this publication was supported by the National Institutes of Health under award numbers DP2CA280626 (JRD), CA207729 (JRD), GM132055 (CMJ), and GM119512 (DTL). Supported in part by the Translational Science Laboratory Shared Resource, Hollings Cancer Center, Medical University of South Carolina (P30 CA138313). The MUSC Proteogenomics Facility was used and is supported by GM103499 and MUSC’s Office of the Vice President for Research. Funding bodies had no role in the design of the study and collection, analysis, and interpretation of data and in writing the manuscript. The content is solely the responsibility of the authors and does not necessarily represent the official views of the National Institutes of Health or other funders.

## Author contributions statement

All authors were involved in writing the paper and had final approval of the submitted versions.

### Abbreviations

CCLE: Cancer cell line encyclopedia
CNA: Copy number alteration
CNV: Copy number variant
FTE: Fallopian tube epithelium
GEMM: Genetically engineered mouse model
H&E: Hematoxylin and eosin
HA: Hemagglutinin
HGSOC: High-grade serous ovarian cancer
HMM: Hidden Markov model
HR: Hazard ratio
NGS: Next-generation sequencing
OSE: Ovarian surface epithelium
OvTrpMyc: Transgenic mouse lineTrp53^em1Jdel_Tg(Ovgp1-Trp53*R270H-Myc)^
STIC: Serous tubal intraepithelial carcinoma
TPM: Transcripts per million reads

