## Supplemental Figures for "MYC is sufficient to generate mid-life high-grade serous ovarian and uterine serous carcinomas in a p53-R270H mouse model"

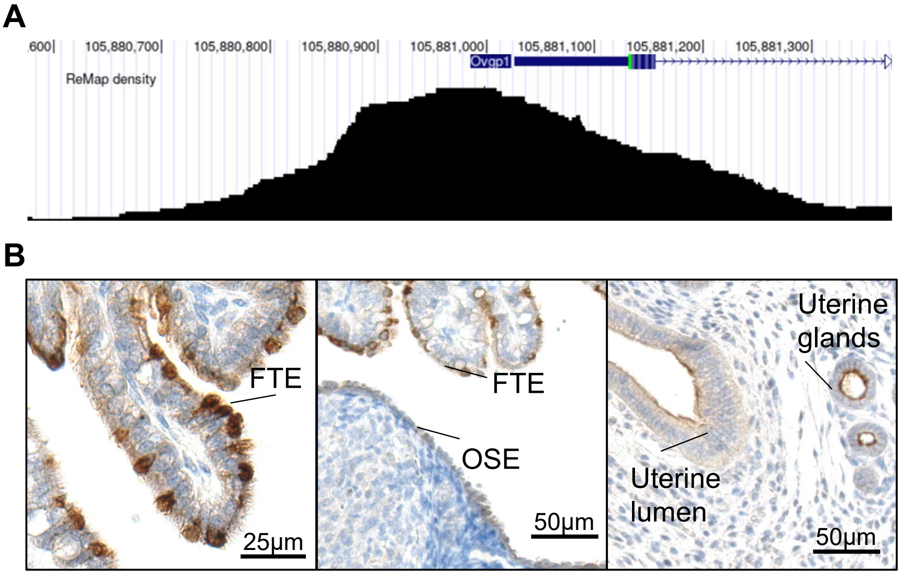


**Figure S1: *Ovgp1* transgene expression**. (A) UCSC Genome Browser display of the murine *Ovgp1* promoter with its regulatory peak, as annotated in the ReMap ChIP-seq database. (B) 3xHA tag transgene immunohistochemical staining at 10 weeks of age in an OvTrpMyc female.

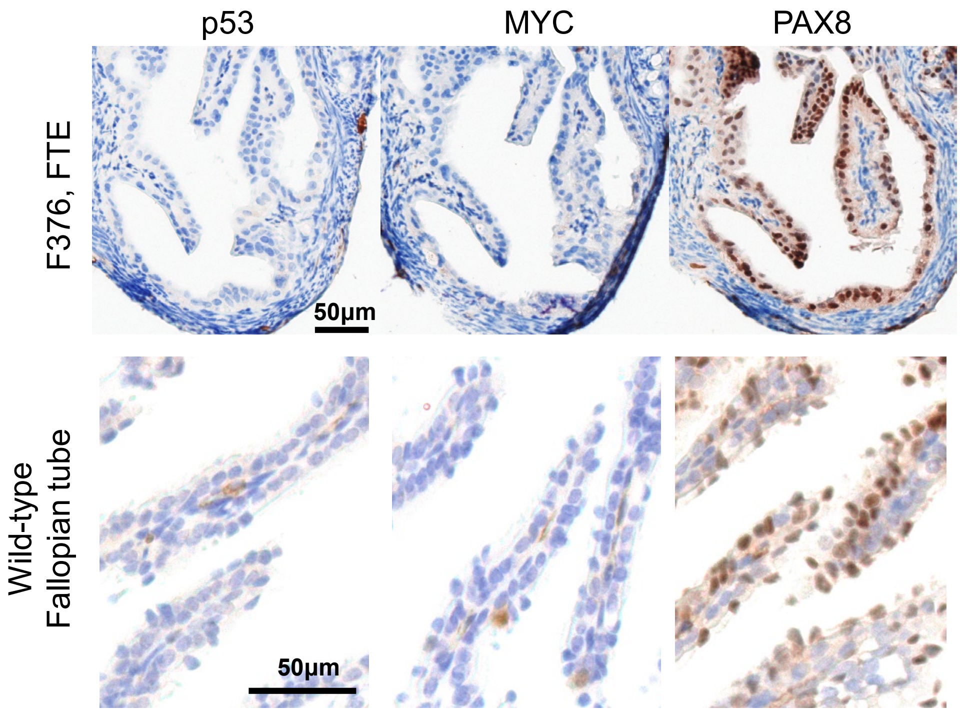


**Figure S2: Absence of p53 and MYC staining in mouse with uterine epithelial expression of p53 and MYC**. Staining of the FTE of OvTrpMyc mouse F376, associated with Fig. 5. Lower panel compares to wild-type mouse staining, at 12-months of age

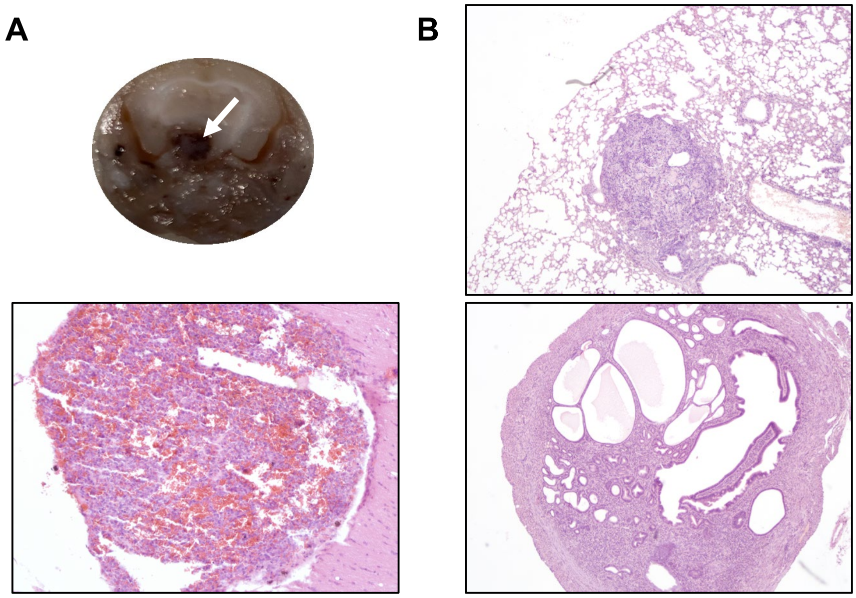


**Figure S3: Necropsy of mice lacking macroscopic disease**. OvTrpMyc mouse were evaluated by necropsy when macroscopic tumors were not immediately observed. **(A)** Dissected brain (top panel) and H&E section of brain tumor (bottom panel). **(B)** Lung metastasis (top panel) and a normal phenotype of some aged mice: cystic uterus (bottom panel).

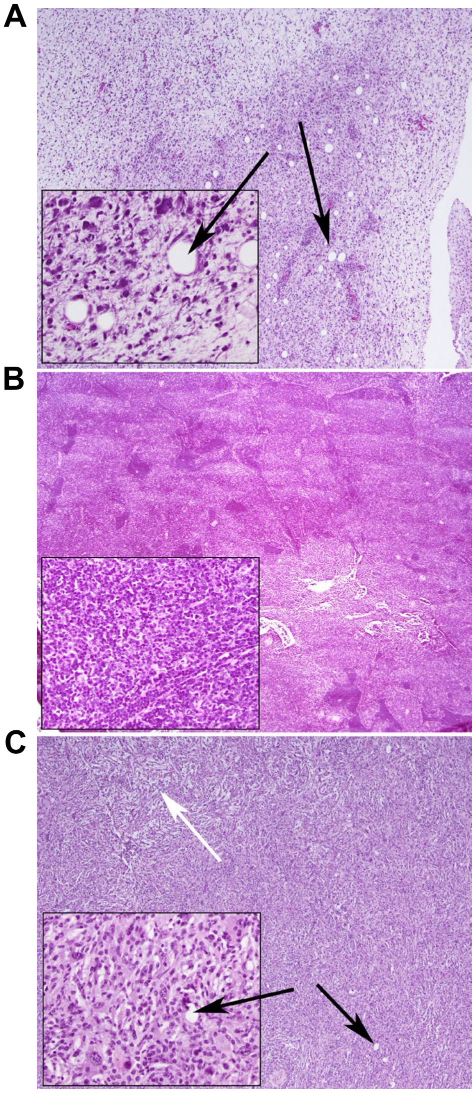


**Figure S4: Other malignancies**. (**A**) Myxoid neoplasm with markedly pleomorphic cells with associated vacuoles, marked by the black arrows. (**B**) Small round blue cell malignancy with relatively uniform nuclei growing in a sheeted growth pattern. (**C**) Tumor characteristics include spindle cell malignancy with scattered giant cells, storiform architecture, marked by the white arrow. Moderately pleomorphic with scattered vacuoles, marked by black arrows.
